# Generating Correlated Data for Omics Simulation

**DOI:** 10.1101/2025.01.31.634335

**Authors:** Jianing Yang, Gregory R. Grant, Thomas G. Brooks

## Abstract

Simulation of realistic omics data is a key input for benchmarking studies that help users obtain optimal computational pipelines. Omics data involves large numbers of measured features on each samples and these measures are generally correlated with each other. However, simulation too often ignores these correlations, perhaps due to the inconvenience and computational hurdles of doing so. To alleviate this, we describe in detail three approaches for quickly generating omics-scale data with correlated measures which mimic real data sets. These approaches all are based on a Gaussian copula approach with a covariance matrix that decomposes into a diagonal part and a low-rank part. We use these approaches to demonstrate the importance of including correlation in two benchmarking applications. First, we show that variance of results from the popular DESeq2 method increases when dependence is included. Second, we demonstrate that CYCLOPS, a method for inferring circadian time of collection from transcriptomics, improves in performance when given gene-gene dependencies in some circumstances. We provide an R package, dependentsimr, that has efficient implementations of these methods and can generate dependent data with arbitrary distributions, including discrete (binary, ordered categorical, Poisson, negative binomial), continuous (normal), or with an empirical distribution.

## Introduction

Omics data typically has far fewer samples than measurements per sample. This creates dual challenges in generating realistic simulated data for the purposes of benchmarking. First, there isn’t enough data to be able to compute a dependence structure (e.g., a full-rank correlation matrix). Second, generating omics-scale data with a specified correlation matrix is slow due to the typical *O*(*n*^3^) nature of these algorithms, where *n* is the number of measurements per sample. Moreover, there is a lack of practical guidance on how to generate simulated data with realistic dependence. These often mean that simulators assume independence of the measurements, which does not reflect reality.

Here, we give an introduction to the theory and practice of generating dependent data and describe three related solutions which all offer good performance and ease-of-use even for large omics-scale problems. This expands off a discussion we originally wrote as part of a larger discussion on best practices in omics benchmarking (Brooks et al. 2024). Our goal here is to produce guidelines that show that generating correlated data does not have to be onerous and instead should be considered a baseline requirement when simulating data.

We present three methods that operate by inferring a covariance matrix that decomposes into a diagonal part and a low-rank part. Using a Gaussian copula (Nelsen 1998) approach (also referred to as NORTA, for “normal to anything” (Cario and Nelson 1997)), the marginal (univariate) distributions can have realistic forms. These solutions operate by taking a real dataset and mimicking it. For ease of use, we implement this in an R package which supports normal, Poisson, DESeq2-based (negative binomial with sample-specific size factors), and empirical (for ordinal data) marginal distributions.

We implemented three different strategies for determining the diagonal and low-rank parts of the covariance matrix. First, the ‘PCA’ method uses principal component analysis (PCA) and picks the low-rank part such that the simulated data has the same variance in the top *k* PCA components of the reference dataset. Second, the ‘spiked Wishart’ method fits *k* components such that simulations with the same number of samples as the reference dataset will have, on average, the same PCA component variances as the reference. Unlike ‘PCA’, these variances are computed with resepect to the simulated data’s own PCA and not using the PCA weights of the reference dataset. Third, the ‘corpcor’ method uses the popular R library `corpcor` (Schäfer and Strimmer 2005; Opgen-Rhein and Strimmer 2007), which implements a James-Stein type shrinkage estimator for the covariance matrix as a linear interpolation of the sample covariance matrix and a diagonal matrix. No method exactly captures the input data, indicating room for future research, but all improve upon the common approach of assuming independence.

We show two applications which demonstrate the effects of including dependence of measurements in simulated data when benchmarking computational pipelines. First, we simulate RNA-seq data with differential expression between two conditions. Using DESeq2 to determine the differentially expressed genes, we found that dependence had little impact on the accuracy of reported *p*-values but increased the variance of those estimates. Second, we simulated a time series of RNA-seq data points and used the CYCLOPS method (Anafi et al. 2017) to infer collection time from the RNA-seq data, without time labels. Depending upon settings used, performance of CYCLOPS dependent substantially on the dependence structure of the data, and surprisingly showed worst performance when given data with independent genes.

## Results

Assume that we have a reference dataset *X* given by an *p* × *n* data matrix of *p* features measured in each of *n* independent samples. We want to capture correlations between the *p* features, which could represent gene expressions, protein abundances, or other measured values. We refer to these features as genes for simplicity. Our goal is to generate simulated data with the same *p* genes, the same marginal distributions of each gene as in *X* and realistic gene-gene dependence.

### Multivariate normal distribution

We first discuss the simplest case, where our data set is multivariate normally distributed. The distribution *N*(*μ*, Σ) is the multivariate normal distribution with mean vector *μ* and covariance matrix Σ. Here *μ* is a *p* × 1 column vector and Σ is a *p* × *p* matrix. In order for this to work, Σ must be symmetric and positive semi-definite, meaning that all of its eigenvalues are non-negative. This matrix is conceptually simple, since Σ_*ij*_ gives the the covariance of *x*_*i*_ and *x*_*j*_ when *x* ∼ *N*(*μ*, Σ). More specifically, this is the population covariance matrix of *N*(*μ*, Σ), which does not generally equal the sample covariance matrix. Indeed, if *X* is *n* samples from *N*(*μ*, Σ), then 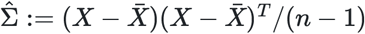is the sample covariance matrix where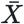is the average of the *n* samples. For large *n*,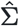 will closely approximate Σ, but we care primarily about the situation where *n* is small. In particular, 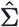 is at most a rank *n* − 1 matrix while Σ could be up to rank *p*. This means that 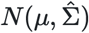 and *N*(*μ*, Σ) are quite different distributions: every sample from 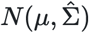 is contained in an *n* − 1 dimensional plane. If *n* is small, then this is very unlike real data, which typically is close to *p* dimensional.

The difficulty then is that we only know 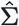from our reference data set, but we need to choose a Σ with which we can simulate data and the obvious choice of 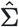 is inadequate. The most common choice is to assume Σ is a diagonal matrix. This is the situation we want to avoid where the generated data is independent: *x*_*i*_ and *x*_*j*_ have zero covariance unless *i* = *j*. However, this has some nice properties, such as being simple and fast to simulate (just generate univariate normal data for each variable).

We describe three alternative approaches in Methods of how to choose a Σ that will produce data similar to the input data. All three of these rely on a specific form of Σ, namely that

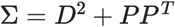

where *D* is a *p* × *p* diagonal matrix and *P* is *p* × *k* for some *k* ≪ *p*. This is a combination of an independent part (the diagonal matrix) and a low-rank part (*PP* ^*T*^). The low-rank part is also simple to generate data for. If *x* is a vector of *k* independent univariate standard normal values, then *Px* ∼ *N*(0, *PP* ^*T*^). The choice of specific *D* and *P* matrices is more in-depth and we leave the details for the Methods section. We have three alternative means for doing so, which we refer to as the PCA, spiked Wishart, and corpcor methods.

Lastly, we emphasize that multivariate normal distributions do not capture all, or even most, types of possible dependence. Indeed, we see this even in the 2-dimensional case where it is well known that correlation describes only a linear relationship between two variables while in reality they may have much more complex relations. In higher dimensions, the problem is only worse. So any method based off multivariate normal distributions are making large assumptions about distribution. However, it is necessary to make some assumption like this. In the next section, though, we see that “normal” part is actually not a large obstacle.

### Gaussian copula

Building on the multivariate normal distribution, a popular approach to describe dependence in a high-dimensional settings is called the Gaussian copula approach. The idea of this approach is that by normalizing and later reversing the normalization, data that does not fit a normal distribution can still have its dependence structure described using a multivariate normal distribution. This allows the marginal (i.e., univariate) distributions of each genes to be specified separately from the dependence between genes. This operates first by normalizing each gene by fitting a distribution (such as a normal distribution, Poisson, negative binomial, or other form), and then applying the fit cumulative distribution function (CDF) to the observed values. Finally, those are fed to a standard normal distribution’s inverse CDF to obtain values that are approximately normally distributed. These values are then used to compute a covariance matrix Σ and the data is assumed to follow a multivariate normal distribution in *p* dimensions with that covariance matrix.

Here, we describe the approach using the form of covariance matrix Σ = *D*^2^ + *PP* ^*T*^ as above. Once data is obtained *Z* ∼ *N*(0, Σ), then one can undo the normalization process to obtain data with the same marginal distributions as the fit marginal distributions but with dependence determined by Σ. We describe this in detail:

1. Fit marginal distributions to each feature in *X* to determine CDFs *F*_*i*_ for each feature.
2. Transform *X* to normalized values by *Z*_*ij*_ = Φ^−1^(*F*_*i*_(*X*_*ij*_)) where Φ is the CDF of the standard normal distribution.
3. Compute *D, U*, *W* matrices from *X* by one of three methods (see Methods).
4. Generate *k* i.i.d. standard normally distributed values *u* and *p* i.i.d standard normally distributed values *v*.
5. Set *Z* ^′^ = *UWu* + *Dv*.
6. Output the vector *X* ^′^ where 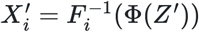.

The generated data *Z* ^′^ has covariance matrix Σ = *D*^2^ + *UWU* ^*T*^ = *D*^2^ + *PP* ^*T*^, where 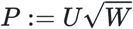. Moreover, we require that Σ satisfies that 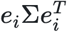 is approximately 1. That guarantees that the output *X* ^′^ has each entry with the same marginal distributions *F*_*i*_ as was originally fit and inherits gene-gene dependence from *Z* ^′^. This method is computationally efficient, taking hardly any more time or memory than simulations without dependence.

### Comparison to real data

To compare the three simulation methods with a real data set, we chose as a references data set 12 mouse cortex RNA-seq samples from accession GSE151923 (Wang et al. 2022). We then simulated data mimicking this reference using all three simulation methods (PCA, spiked Wishart, and corpcor) as well as a simulation with independent genes. We repeated the simulations, each of 12 samples, a total of 8 times to estimate variance. For the PCA method, we used *k* = 2 dimensions and for the spiked Wishart, *k* = 11. The coprcor method always uses the full data matrix, analogous to *k* = 11. Note that PCA method must use a rank *k* < 11 in order to generate full-rank data, see Methods, so these parameters are not directly comparable across methods.

Simulated data captures the genes’ mean and variance accurately (Figure 1 a-b). Next, we compared to the real data set when projected onto the top two principal components of the real data set (Figure 1 c). The simulations with dependence are distributed around the entire space like the real data, but the independent simulations have unrealistically low variance in these components, clustering tightly around the origin.

**Figure 1.**
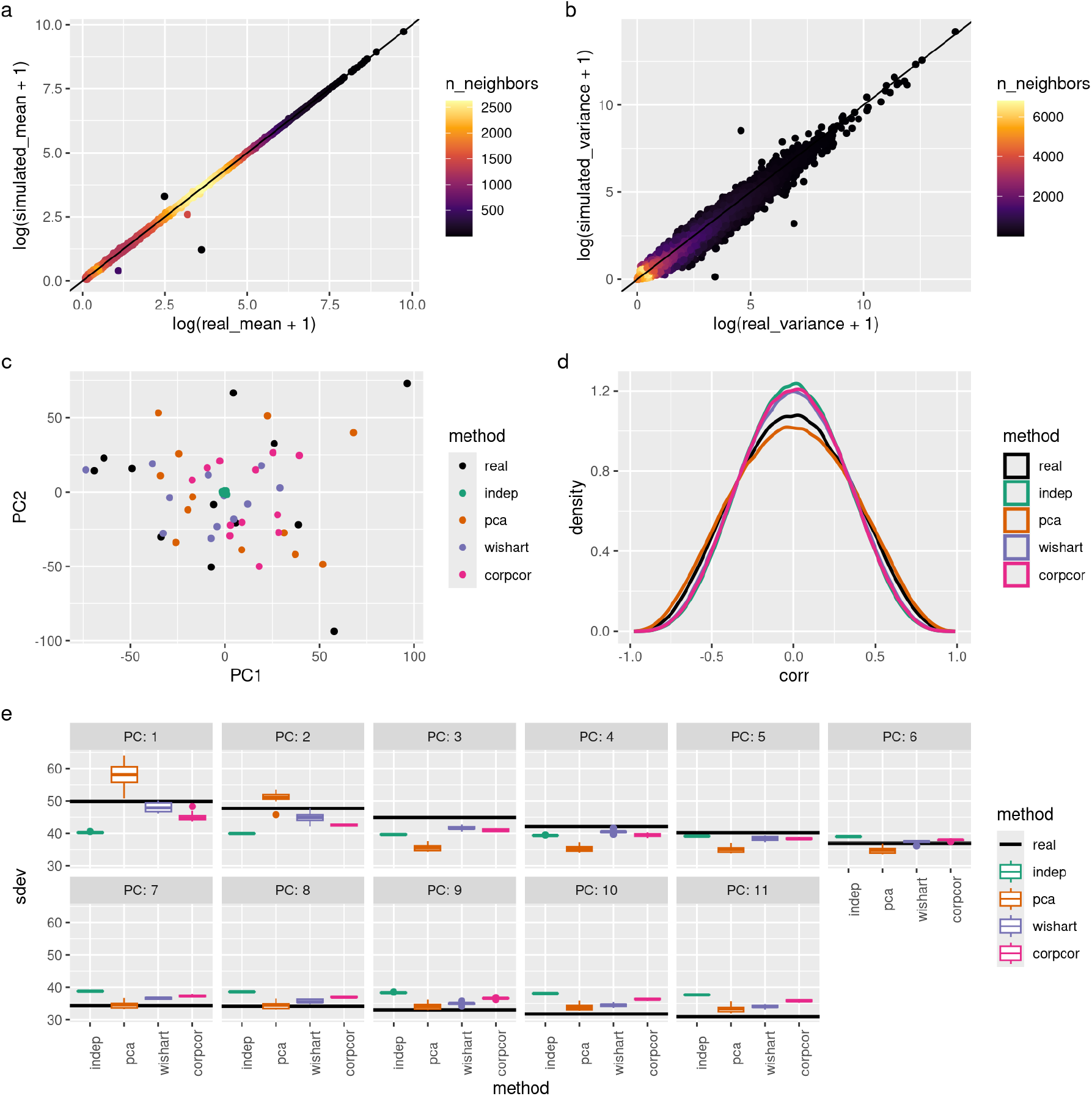
Comparison to real data run on a mouse cortex data set from GSE151923. (a-b) Comparison of gene (a) mean expression and (b) variance, log-scaled in real and PCA simulated data. The line of equality is marked in black. Points are colored according to the density of points in their region. Wishart and corpcor methods give similar results (not shown). (C)Quantile-quantile plot comparing correlation values of gene pairs from real data and simulated data (both with and without dependence). Genes with at least 300 reads were used. Values on the diagonal line indicate a match between the simulated and real data sets. (d) Projections onto the top two principal components of the real data set for both real and simulated data. All 8 simulations (96 samples for each simulation) shown. (e) Principal component analysis was performed on all data sets and the variance captured by the top components is shown. Unlike (d), these components were fit from each data set considered separately instead of reusing the weights from the real data.

Then, we computed the gene-gene correlation on pairs of high-expressed genes (at least 300 mean reads). The simulation with independence showed the least levels of gene-gene correlations (Figure 1 d). However, the PCA method overshot the reference data set and the spiked Wishart and corpcor methods only slightly improved upon the independent simulation.

Lastly, we compared the variances of principal components on each data set (Figure 1 e). These were computed separately for each data set, unlike (Figure 1 c) which used the reference data set’s PCA weights for all data sets. The independent data has much lower variance than the real data set in the top four principal components. The spiked Wishart method comes closest to the real data set, as it optimizes for fitting these values. Surprisingly, the corpcor method performs only somewhat better than the independent method. The PCA method puts a large amount of variance into the first two components (due to using *k* = 2) and then undershoots the other components.

### DESeq2 application

We benchmarked DESeq2 (Love, Huber, and Anders 2014), a popular differential expression analysis tool, using data sets simulated with dependence and ones simulated without dependence to compare its performances on both. DESeq2 presents an interesting case because several aspects of it assume independence of genes and so may be adversely affected by gene-gene dependence. First, the independent filtering step (Bourgon, Gentleman, and Huber 2010) assumes independence but has been reported to be robust to typical gene-gene dependence. Relatedly, the false discovery rate (FDR) (Benjamini and Hochberg 1995) allows only certain forms of dependence. Lastly, DESeq2’s empirical Bayes steps could possibly be affected by gene dependence.

We used a fly whole body RNA-Seq data set GSE81142 and selected samples of male flies without treatment and after at least 2 hours of feeding to simulate 5 “control” samples. We then randomly selected 5% of the genes to be differentially expressed, with absolute log_2_ fold change uniformly distributed between 0.2 and 2.0, either up or down regulated chosen randomly, and simulated 5 “experimental” samples. This was repeated 20 times for each of four dependence conditions (independent, PCA, Wishart, and corpcor).

Finally, we ran DESeq2 on each simulated 5 vs 5 experiment and compared the output FDR with the true percentages of genes that are differential expressed (Figure 2 a-d). We observed that DESeq2 is anti-conservative on all data sets, with similar mean true FDRs for each estimated FDR cutoff. However, there was a greater variance in the performance of DESeq2 on the data sets simulated with dependence, indicating that it performs less consistently on data sets with gene-gene dependence, as in real data sets.

**Figure 2.**
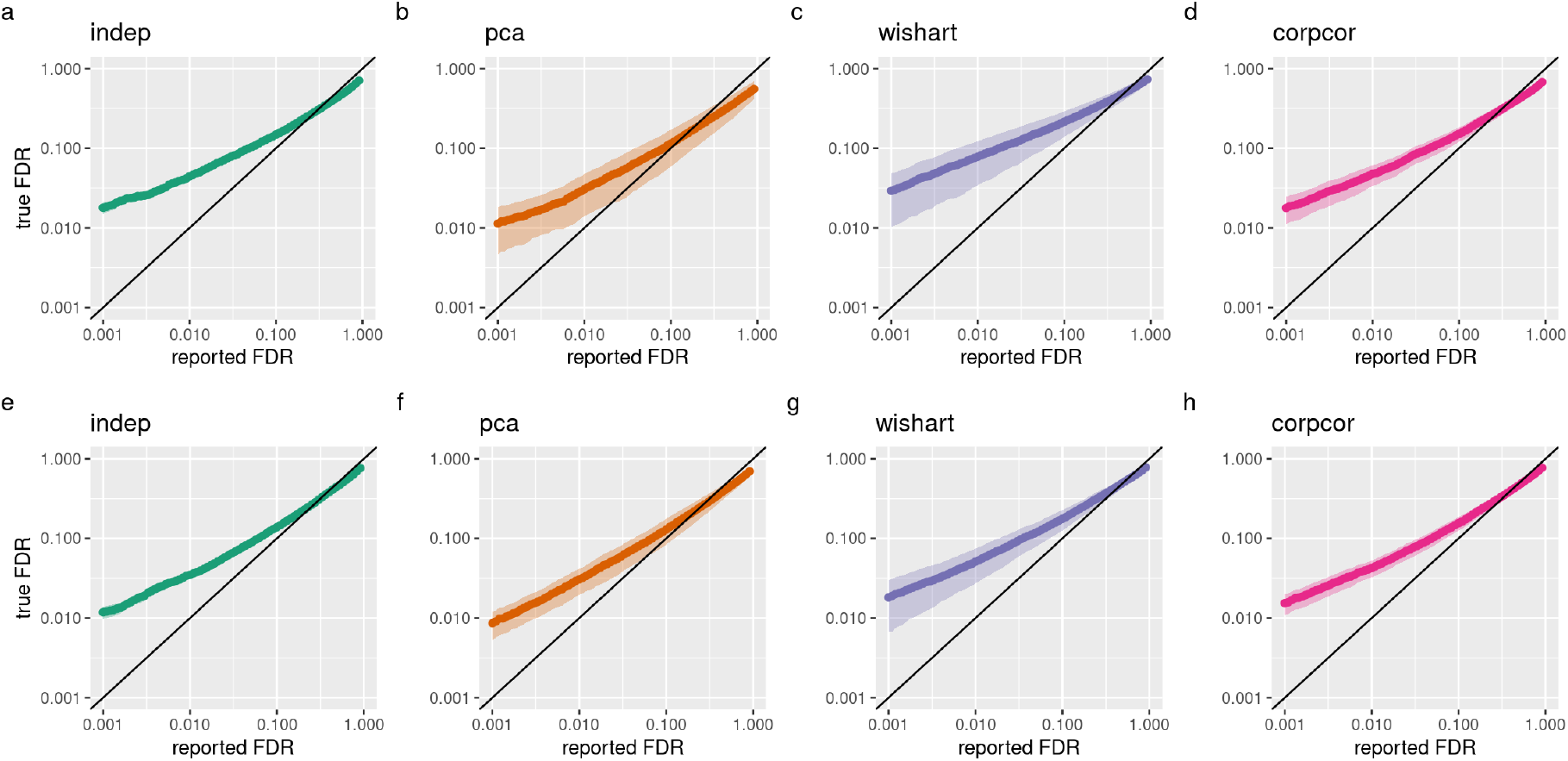
Performance of DESeq2 on simulated datasets. (a-d) Comparison of true false discovery proportions and DESeq2 reported False Discovery Rates, plotted on a log scale, for data sets simulated from the fly whole body data set (GSE81142), (a) without dependence, (b) using PCA, (c) using Wishart and (d) using corpcor. Diagonal line represents perfect estimation of FDR. (e-h) Comparison of true false discovery proportions and DESeq2 reported FDR for data sets simulated from the mouse cortex data set (GSE151923), (e) without dependence, (f) using PCA, (g) using Wishart and (h) using corpcor.

To demonstrate the application of our simulation method for another organism, we also simulated data sets using mouse cortex data set GSE151923 (Wang et al. 2022) and selected samples from male mice. We then simulated differential expression experiments as above and observed a similar result (Figure 2 e-f) as for the fly whole body data sets.

### CYCLOPS application

We next used our simulation method to benchmark CYCLOPS (Anafi et al. 2017), which infers relative times for a set of unlabeled samples using an autoencoder to identify circular structures. We chose a mouse cortex time series data set GSE151565, which contains a total of 77 samples every 3 hours, for 36 hours. We computed the dependence structure of the genes as well as the variances of marginal distributions using the 12 time point 0 samples and computed the means of gene expressions at each time point. We then used these to simulate 20 time series data sets for each of the independent, PCA, Wishart, and corpcor simulation methods.

We ran CYCLOPS on each data set with a list of cyclic mouse genes (from (Zhang et al. 2014), JTK p-value < 0.05), which yielded an estimated relative time for each sample. We evaluated CYCLOPS’ performance compared to true circadian time using the circular correlation (Fisher and Lee 1983), defined as follows:

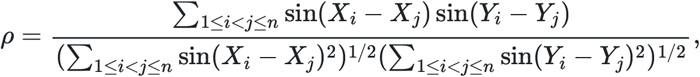

where *n* is the number of samples, *X*_*i*_ and *Y*_*i*_ are the true time and CYCLOPS-estimated time, respectively, for the *i*-th sample. *ρ* has value between −1 and 1, and a |*ρ*| close to 1 indicates accurate predictions by CYCLOPS.

By default, CYCLOPS performs dimension reduction so that each dimension (called an “eigengene”) contains at least 3% of the total variance. We found that CYCLOPS performance depended significantly on this parameter, with the default producing good performance across all simulations. However, when dropping CYCLOPS to require just 2% variance in each eigengene, we found that its performance depends significantly on the dependence structure of the simulated time series data (Figure 3 a). At that setting, CYCLOPS performance is much higher in the PCA method and moderately improved in Wishart method, compared to the independent method. This difference is likely driven by the difference in the number of eigengenes used (Figure 3 b), which is a measure of how much dependence is present in the data set. This demonstrates that the correlational structure of the transcriptome can have a major impact on performance.

**Figure 3.**
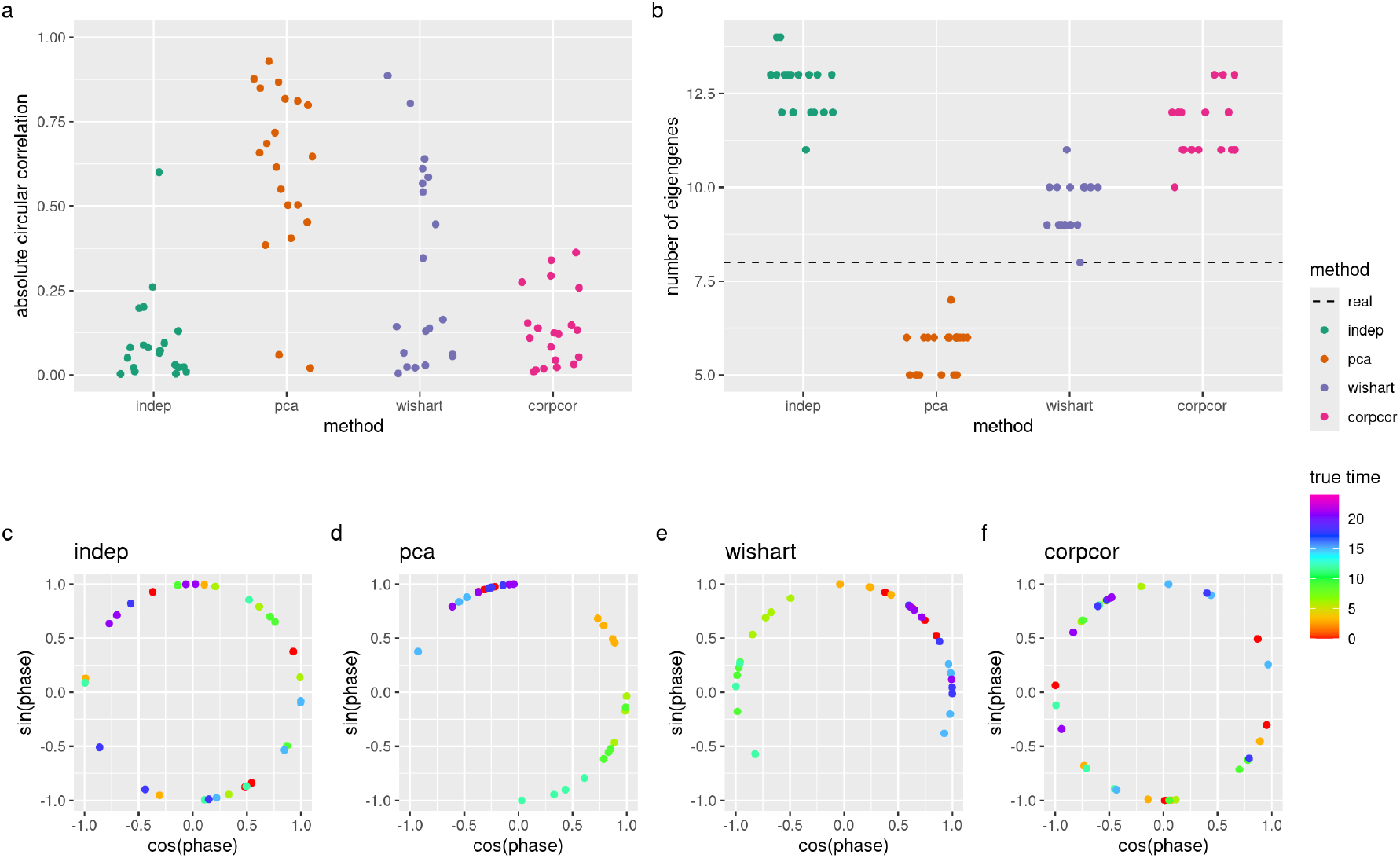
Performance of CYCLOPS on simulated time series data sets based on mouse cortex data set (GSE151565), using eigengenes of at least 2% variance. (a) Absolute circular correlations between true phases and CYCLOPS estimated phases on the simulated data sets. (b) The number of eigengenes used by CYCLOPS; the dotted line indicates the number of eigengenes used by CYCLOPS on the real data (5). (c-f) Examples of CYCLOPS estimated phases on the simulated data sets. CYCLOPS shows good performance when it separates out points by color (true circadian time).

## Methods

Below, we describe the three methods for selecting the components of the covariance matrix Σ := *D*^2^ + *PP* ^*T*^.

### PCA method

The first of our three methods attempts to match the top *k* PCA components of the normalized reference dataset *Z*. Specifically, let *u*_1_, …, *u*_*k*_ be the left singular vectors of *Z* with *λ*_1_, …, *λ*_*k*_ the corresponding top *k* signular values. This method computes Σ such that 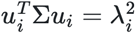, i.e. that the variance in the direction of *u*_*i*_ exactly matches of the reference dataset’s variance in that same direction. One solution is to use the sample covariance matrix, but that is not full rank and would match for all *i* ≤ *n* instead of just *i* ≤ *k*.

Instead, we use the following:

1. Compute 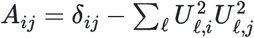and 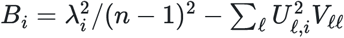 where *δ*_*ij*_ is the Kronecker delta and *V* = *Z*^*T*^ *Z*/(*n* − 1) is the covariance matrix of *Z*.
2. Solve *Aw* = *B* and set *W* to be the diagonal matrix with *w* along its diagonal.
3. Set *U* to be the *p* × *k* matrix with columns *u*_*i*_.
4. Set *D* to be the diagonal matrix with 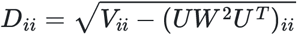, which is the remaining variance.

Steps 1 and 2 give that 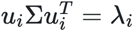, …, *k*. Step 4 ensures that 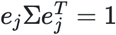for *j* = 1, …, *p*.

### Spiked Wishart method

The second method also makes use of PCA but has a different objective. If *n* samples are drawn from *N*(0, Σ) then we want the variances of their PCA components to match those of the reference dataset. Specifically, let *λ*_1_, …, *λ*_*n*−1_be the *n* − 1 non-zero singular values of *Z* (*λ*_*n*_ is always approximately zero due to the normalization procedure), and let 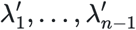 be the singular values of *Z* ^′^ where *Z* ^′^ has *n* – 1 columns each iid *N*(0, Σ). Then we want to choose Σ such that 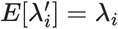 for each *i*, where *E*[*Y*] denotes the expectation of the random variable *Y*.

Since the distribution of the *λ*^′^ does not have a known analytic solution, we approximate this situation with the spiked Wishart distribution. The rank *k* − 1 Wishart distribution is that of the sample covariance matrix of *Y* where the *n* columns of *Y* are iid *N*(0, Σ). The spiked Wishart is the special case where Σ has *k* arbitrary eigenvalues and the remaining are all equal to a constant. Note that the singular values of *Y* are the square roots of the eigenvalues of its sample covariance matrix. While our case has Σ non-diagonal, Σ may be diagonalized by orthogonal rotations due the sepctral theorem, and orthogonal rotations do not change the singular values of *Y*. Therefore, the distribution of singular values is not affected by the assumption that Σ is diagonal. Moreover, for the form Σ = *D*^2^ + *UWU* ^*T*^ where *p* is very large, each column of *U* is typically very close to orthogonal to any *e*_*i*_, a standard basis vector. Therefore, when *D* = *c*^2^*I* for some constant *c*, we can approximate Σ as having *k* arbitrary eigenvalues from *W* and *n* remaining eigenvalues all equal to *c* corresponding to *D*. This is a spiked Wishart distribution.

However, the spiked Wishart distribution also has no known analytic solution for the distribution of its eigenvalues either. Therefore, we use an efficient sampling and stochastic gradient descent method that we recently described (Brooks 2024). Since the dataset has been normalized, *c* will be close to one and 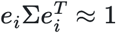.

Specifically, we do:

1. Set *U* to be the *p* × *k* matrix with columns *u*_*i*_, the left singular vectors of *Z*.
2. Compute *w*_1_, …, *w*_*k*_ and *c* by stochastic gradient descent minimizing 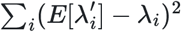 for Σ diagonal with entries *w*_1_, …, *w*_*k*_, *c*^2^, …, *c*^2^ (Brooks 2024).
3. Set *W* diagonal with the entries *w*_1_, …, *w*_*k*_.
4. Set *D* = *cI*

### Corpcor method

The `corpcor` package (Schäfer and Strimmer 2005; Opgen-Rhein and Strimmer 2007) computes a James-Stein type shrinkage estimator for the covariance matrix. For large *p*, this greatly improves the estimate of the covariance matrix by introducing a little bias towards zero correlations and equal variances of genes. It computes optimal values of *λ*_1_, and *λ*_2_, its two regularization coefficients. It then uses *λ*_1_ to linearly interpolate the sample covariance matrix towards the identity matrix *I* and *λ*_2_ to interpolate the vector of variances towards the median variance value. Since the sample covariance matrix is rank at most *n*, we again obtain a matrix of the form Σ = *D*^2^ + *UWU* ^*T*^.

This algorithm is:

1. Compute the *λ*_1_ and *λ*_2_ values from corpcor::estimate.lambda and corpcor::estimate.lambda.var functions on *Z*, respectively.
2. Set *D* to be diagonal with 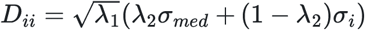where *σ*_*i*_ is the standard deviation of the *Z*_*i*_· and *σ*_*med*_ is the median of the *σ*_*i*_.
3. Set *U* to be 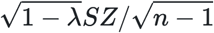 where *S* is the diagonal matrix with *S*_*ii*_ = *λ*_2_*σ*_*med*_/*σ*_*i*_ + (1 − *λ*_2_).
4. Set *W* to the identity.

## Discussion

We described the well-known Gaussian copula approach and recommended a specific form of covariance matrix which is well tailored to omics data simulation. We developed three methods using this form of covariance matrix which can be used to mimic a reference data set for simulation. All of these methods use a multivariate normal distribution as an intermediate step and therefore substantially restrict the kinds of dependence that can be simulated. However, when operating in a high-dimensional space some simplification is likely required.

To encourage adoption of dependence in simulated omics data, we developed `dependentsimr`, an R package that generates omics-scale data with realistic correlation. This implementation is efficient and simple, requiring just two lines of code to fit a model to a reference data set and then simulate data from it. We demonstrated this package on RNA-seq data, using the DESeq2 method to fit negative binomial marginal distributions. However, this package is actually quite general and supports normal, Poisson, negative binomial, and arbitrary ordered discrete distributions using the empirical CDF. Moreover, it can support multi-modal data such as is increasingly common in multi-omics.

We demonstrated the importance of including gene-gene dependence in simulated data by two application benchmarks. In the first, DESeq2 results were substantially more variable when simulating with gene-gene dependence. In the second, CYCLOPS performance in estimating circadian phases depends upon gene-gene dependence, was sensitive to dependence structure of the data.

Our comparisons to a real dataset show that none of our three methods are able to exactly capture all aspects of the real dataset. In particular, the gene-gene correlations were too high in the PCA method and too low in the spiked Wishart and corpcor methods. Surprisingly, the spiked Wishart and corpcor methods improved in this metric only slightly compared to the simulations with independent genes. These observations demonstrate that there is room for future improvements over independent data in these techniques, possibly incorporating more recent developments in copulae (Größer and Okhrin 2022).

Possibly, this could demonstrate the limitations of methods based on the multivariate normal distribution or of the low-rank approximation used by all three of our methods. Nonetheless, these methods represent significant improvements by other metrics and we recommend the inclusion of some dependence in nearly every simulated omics dataset.

### Alternatives

We highlight some alternative approaches and software packages that have been taken below.

The R package SPsimSeq (Assefa, Vandesompele, and Thas 2020) provides a dedicated RNA-seq and single-cell RNA-seq simulator using a Gaussian copula approach to simulate gene dependence. In contrast to this package, it uses WGCNA to determine the correlation matrix, which is a gene network approach. However, this method takes significant computational resources. Indeed, the `SPsimSeq` paper generated data for just 5000 genes based on a randomly sampled 5000 gene subset of the RNA-seq data and our attempts to use `SPsimSeq` to generate a full sample exhausted the memory of a 24GB computer. In contrast, our method runs in seconds to generate a 40,000 gene samples on the same computer. `SPsimSeq` is more specialized and full-featured for RNA-seq simulation, providing, for example, native differential expression (DE) options. In comparison, our `dependentsimr` package requires manually setting marginal expression values to inject DE, but also supports other marginal distributions for situations outside of RNA-seq.

The scDesign2 simulator (Sun et al. 2021) for single-cell RNA-seq also uses Gaussian copula and, like our method, uses the approach of estimating the correlation matrix from the normalized dataset. However, it limits this correlation matrix to top-expressed genes. Since correlation is most discernible in high-expressed genes, this approach is reasonable but requires making certain arbitrary cutoffs that our methods avoid.

Other Gaussian copula-based R packages that may be applicable, at least for datasets with smaller numbers of features, include `bindata`, `GenOrd`, and `SimMultiCorrData`, the last of these being the most comprehensive. The `bigsimr` package provides faster implementations of these methods to scale up to omics-level data. However, even this is computationally demanding; their paper references generating 20,000-dimensional vectors in “under an hour” using 16 threads. The `copula` package provides even more flexible dependence options through use of copulas. All of these packages provide more flexibility in specifying dependence than our package, which can only mimic existing datasets, and therefore the longer run-times may be unavoidable for use cases where researchers need to parameterize the dependence structure.

## Data availability

Source code for all simulations and figures in this plot is available at github.com/itmat/dependent_sim_paper/. Source code for the `dependentsimr` package is available at github.com/tgbrooks/dependent_sim. All data used is available with accession numbers GSE151923, GSE81142, GSE151565.

## References

Anafi, Ron C, Lauren J Francey, John B Hogenesch, and Junhyong Kim. 2017. “CYCLOPS Reveals Human Transcriptional Rhythms in Health and Disease.” Proc. Natl. Acad. Sci. U. S. A. 114 (20): 5312–17.

Assefa, Alemu Takele, Jo Vandesompele, and Olivier Thas. 2020. “SPsimSeq: Semi-Parametric Simulation of Bulk and Single-Cell RNA-sequencing Data.” Bioinformatics 36 (10): 3276–78.

Benjamini, Yoav, and Yosef Hochberg. 1995. “Controlling the False Discovery Rate: A Practical and Powerful Approach to Multiple Testing.” Journal of the Royal Statistical Society. Series B (Methodological) 57 (1): 289–300. http://www.jstor.org/stable/2346101.

Bourgon, Richard, Robert Gentleman, and Wolfgang Huber. 2010. “Independent Filtering Increases Detection Power for High-Throughput Experiments.” Proc. Natl. Acad. Sci. U. S. A. 107 (21): 9546–51.

Brooks, Thomas G. 2024. “Sampling Spiked Wishart Eigenvalues.” https://arxiv.org/abs/2410.05280.

Brooks, Thomas G., Nicholas F. Lahens, Antonijo Mrčela, and Gregory R. Grant. 2024. “Challenges and Best Practices in Omics Benchmarking.” Nature Reviews Genetics 25 (5): 326–39. 10.1038/s41576-023-00679-6.

Cario, Marne C., and Barry L. Nelson. 1997. “Modeling and Generating Random Vectors with Arbitrary Marginal Distributions and Correlation Matrix.”

Fisher, N. I., and A. J. Lee. 1983. “A correlation coefficient for circular data.” Biometrika 70 (2): 327–32. 10.1093/biomet/70.2.327.

Größer, Joshua, and Ostap Okhrin. 2022. “Copulae: An Overview and Recent Developments.” WIREs Computational Statistics 14 (3): e1557. 10.1002/wics.1557.

Love, Michael I, Wolfgang Huber, and Simon Anders. 2014. “Moderated Estimation of Fold Change and Dispersion for RNA-seq Data with DESeq2.” Genome Biol. 15 (12): 550.

Nelsen, Roger. 1998. An Introduction to Copulas. Lecture Notes in Statistics. New York, NY: pnSpringer.

Opgen-Rhein, Rainer, and Korbinian Strimmer. 2007. Statistical Applications in Genetics and Molecular Biology 6 (1). 10.2202/1544-6115.1252.

Schäfer, Juliane, and Korbinian Strimmer. 2005. Statistical Applications in Genetics and Molecular Biology 4 (1). doi:10.2202/1544-6115.1175.

Sun, Tianyi, Dongyuan Song, Wei Vivian Li, and Jingyi Jessica Li. 2021. “scDesign2: A Transparent Simulator That Generates High-Fidelity Single-Cell Gene Expression Count Data with Gene Correlations Captured.” Genome Biol. 22 (1): 163.

Wang, Nan, Peter Langfelder, Matthew Stricos, Lalini Ramanathan, Jeffrey B Richman, Raymond Vaca, Mary Plascencia, et al. 2022. “Mapping Brain Gene Coexpression in Daytime Transcriptomes Unveils Diurnal Molecular Networks and Deciphers Perturbation Gene Signatures.” Neuron 110 (20): 3318–3338.e9.

Zhang, Ray, Nicholas F Lahens, Heather I Ballance, Michael E Hughes, and John B Hogenesch. 2014. “A Circadian Gene Expression Atlas in Mammals: Implications for Biology and Medicine.” Proc. Natl. Acad. Sci. U. S. A. 111 (45): 16219–24.

